# Improving tauG-control of action in PD

**DOI:** 10.1101/143313

**Authors:** Benjaman Schögler, Rachel Polokoff, Gert-Jan Pepping, Jon Perkins, David N Lee

## Abstract

A theory of action control (General Tau Theory) is applied to analyzing normal and abnormal movements in PD; and to designing and testing the efficacy of a sonic aid for PD. A central aspect of the theory, which is supported by experimental evidence across a variety of actions and species, is that the trajectories of competent skilled actions follow a particular temporal pattern, which is described by the mathematical function, tauG. Since tauG-control of actions can be largely deficient in PD, we designed a device that generates whoop-like sounds, where the fundamental frequency of the sound follows the tauG pattern. Our hypothesis was that by listening to these sounds the nervous system of someone with PD might be helped subsequently to self-generate tauG patterns in their nervous system, which might facilitate movement control in different situations. Five adults with PD, and five age-matched controls, took part in the study. They each listened to the sounds under two conditions: (a) experimental - turning a handle for 5 minutes while the sounds were played (b) control - turning the handle without the sounds. Before and after each condition, the tauG-control of lateral body sway while standing was measured (without the sounds playing), using force-plates, on two tasks: (i) keeping the feet down, (ii) lifting the trailing foot. The number of participants out of five, who showed greater ratio improvement following practice with whoop sounds compared to without sounds, was, on each task, high for the PDs compared with the age-matched controls (4 vs 2 or 3). Thus, for the PDs, listening to the tauG whoop-like sounds while performing one action (handle turning) improved subsequent tauG-control on a different task (body-swaying).

## Introduction

Parkinson’s disease (PD) is a neural disorder that predominantly affects a person’s ability to execute purposeful movement (Marsden, 1994). Most common activities, like walking, writing, speaking, looking, reaching, grasping, and holding a limb still can be affected. It is generally considered that disorders in the basal ganglia play a central role in the disease – in particular, that death of dopaminergic neurons in the substantia nigra pars compacta depletes the vital supply of the neurotransmitter, dopamine, to brain cells. Consequently, symptoms of PD are mainly managed through prescribed medication (levadopa) to augment the depleted dopamine supply. However, the positive effects of the treatment are off-set by a reduced response over prolonged treatment and increasing negative side effects, such as dyskinesia, with increased dosage (Fahn, 1996). After several years many patients may experience crippling ‘on/off’ phenomena defined by Marsden as “sudden and unpredictable fluctuations in motor disability unrelated to the timing of levadopa intake” (Marsden, 1994 p.675). It is not clear to what extent increased motor dysfunction and dyskinesias in PD are due to the natural progression of the disease or to prolonged exposure to levadopa therapy (Fahn, 1996). As life expectancy is near normal for patients, the question of when to prescribe levadopa to alleviate the symptoms of PD is a complex one, but on the whole recommendations are for delaying such treatment until absolutely necessary (Marsden, 1994). As a consequence supplementary methods of improving or sustaining mobility are a key concern in both the treatment and management of the disease. However, the precise neural functions that are disabled in Parkinson’s disease, leading to poor control of purposeful movement, has remained a mystery, which means that satisfactory treatment of the disease has remained elusive. In this paper we address the neural basis of Parkinson’s disease, and then develop and test a non-invasive method for alleviating the neuromotor symptoms of the disease.

We start by considering the three basic functions of the nervous system that are normally involved in effective control of purposeful actions. These are *prescribing* the action, *perceptually monitoring* the action, and *performing* the action. Consider, for example, piano playing. The musical actions on the keys must be *prescribed* in the pianist’s nervous system, because it is the pianist who determines what to play and how to play it. The pianist’s nervous system must also *perceptually monitor* the musical actions as they unfold, because external forces (such as sticky keys) can knock actions off their intended course. Finally the pianist’s nervous system has to press the keys appropriately so that the perceptually monitored actions match the prescribed actions.

Given that these three basic functions of the nervous system - prescribing, perceptually monitoring and performing - are the foundation of all purposive actions, the question arises: which of these functions are compromised in PD? The phenomenon of paradoxical movement provides a clue (Martin 1967; Glickstein & Stein, 1991; Majsak *et al*., 1998). Someone suffering from Parkinson’s disease may find it difficult to reach out for a stationary ball, for instance, but is able to catch a moving ball with comparative ease. Or they may experience difficulty walking across a clear floor, yet can step out relatively easily when going up or down stairs. The phenomenon of paradoxical movement suggests that it is the *prescriptive* function of the nervous system that is principally impaired in Parkinson’s disease, rather than the perceptual monitoring or the performing functions, because the latter two functions are clearly operating well when carrying out paradoxical movements, like catching a ball or walking down stairs.

What, precisely, is the prescriptive function of the nervous system that might be impaired in Parkinson’s Disease? According to General Tau Theory (Lee 1998, 2009), which is a general theory of action control inspired by the pioneering theories of Gibson (1966) and Bernstein (1967), one way of prescribing how an action-gap should close is to make the tau of the gap (the time to closure of the gap at the current rate of closure) proportional to a particular tau function, called tauG, which equals the tau of a gap that closes from rest at constant acceleration - as when Newton’s apple fell to the ground. The mathematical formula for tauG, 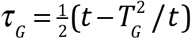, is, appropriately enough, derived from Newton’s equations of motion, where *T_G_* is the time to closure of the *G* gap and time, t, runs from zero to *T_G_*. The tauG-control equation describing the prescribed manner of closure of an action-gap, *X*, is

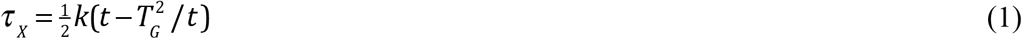

where *τ_X_* is the tau of gap *X*, and *k* is a coupling constant which determines *how* gap *X* is prescribed to close - *gently*, ending with zero velocity, if 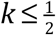; or *forcefully*, ending with positive velocity, if 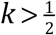.

There is experimental behavioural evidence across a range of activities and species that tauG *τ_G_* is used in controlling purposeful movements (Lee 2009). There is also neural evidence that *tauG* is generated in the nervous system. Electrical activity of tauG form has been recorded in globus pallidus in the basal ganglia of monkeys reaching to targets (Lee *et al*., 2017a). In general, the experiments indicated that globus pallidus is involved in *prescribing* the movement, the subthalamic nucleus is involved in *perceptually monitoring* the movement, and zona incerta is involved in *performing* the movement, by integrating the prescribing activity of globus pallidus and the perceptually monitoring activity of the subthalamic nucleus to power the muscles to perform the movement. In all cases the neural information took the form of the relative-rate-of-change (= 1/*τ*) of the flow of electrochemical energy in neurons, relative to a reference level.

Our behavioural and neural findings, together with the phenomenon of paradoxical movement in PD, suggested that action control might be improved in PD by stimulating the nervous system with tauG patterns. A non-invasive way of doing this is through sound. Musical sound is often a rich source of tauG patterns (Schogler, Pepping & Lee, 2008), which is possibly why people with PD find that music helps them move more easily. To test the efficacy of sonic tauG stimulation *per se* in improving movement control in PD, we designed tauG-patterned whoop-like sounds, where the fundamental frequency moved up an octave in a tauG way.

## Generating tauG-guided sounds

The sounds were generated on a computer. The frequency of a sound was tauG-guided between a start frequency and an end frequency using Eq. (1), viz., 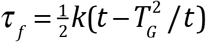, where *f* is the frequency gap to the end frequency, *T_G_* is the duration of the sound and time t runs from zero to *T_G_*. Integrating this equation results in the equation 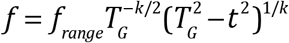, where *f_range_* = end frequency – start frequency. Thus, if a movement gap, *X*, were tau-coupled with coupling constant, *α*, to the frequency-gap, *f* then from Eq. (1), *τ_X_* would equal 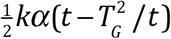 and so gap *X* would be *tauG*-guided. In the present study, an octave frequency range was programmed, starting at 440 Hz (concert A). The duration, *T_G_*, of the sound was 1.2 s, and the coupling constant, *k*, was 0.5. During the experiments the sounds were played to the participants from a laptop computer.

## Participants

Five volunteers with PD and five healthy age-matched volunteer controls took part in the study. The Parkinson volunteers were recruited from the Lothian self-help PD group. The healthy participants either were from the University of Edinburgh or were spouses or friends of the patients. Each participant was informed about the nature of the study and signed a consent form stating that they could terminate their participation in the study at any time. The study was approved by the Ethics Committee of the School of Philosophy Psychology and Language Sciences, University of Edinburgh. A questionnaire was given to the patients to assess their Parkinson symptoms and medication (Table 1). All the patients started trials in the study when at the peak of their medication.

**Table 1.** Details from PD participants’ questionnaire.

| Participant | Gender | Age (yrs.)<br>at testing | Age (yrs.)<br>at diagnosis | Symptoms | Treatment |
| --- | --- | --- | --- | --- | --- |
| 1 | M | 62 | 44 | Difficulty walking, slow gait, tremor, affected right side. | Deep brain stimulation (at 56 yrs.). Ropinirole 3 mg, 3x daily. |
| 2 | M | 65 | 60 | Slow gait, slight tremor, affected left side. | Levodopa. |
| 3 | M | 74 | 49 | Difficulty walking, slow gait, tremor, facial freezing, slow speech, affected right side. | Sinemet Plus (3x daily) |
| 4 | F | 71 | 53 | Difficulty walking, pain in legs, fatigue, tremor in hands, affected right side. | Entacapone (3x daily)<br>Madopar (nightly) |
| 5 | F | 68 | 47 | Slow gait, tiny steps, tremor in hands, affected left side. | Levodopa. |

## Recording body sway

We studied control of lateral body sway because it is integral to maintaining balance when walking, which often poses a problem in PD. During the experiments the participant stood with their feet comfortably apart and swayed from side to side. They stood on two Kistler^™^ forceplates mounted in concrete in the floor of the laboratory, and the (*x*, *y*) coordinates of the centre of pressure of their feet were recorded at 1000 Hz using a Bioware^™^ forceplate program.

## Experimental protocol

Each participant took part in two experimental sessions, spaced about two weeks apart. Each session comprised two parts, and each part comprised three 30s tasks. The tasks are shown in Table 2. ‘Sway side-to-side’ means stand with the feet comfortably apart on the two forceplates and sway from side to side. ‘Keep feet down’ is illustrated in Fig. 1a. ‘Lift trailing foot’ is illustrated in Fig. 1b. ‘Turn wheel’ means rotate a 40 cm diameter wheel mounted at a comfortable height on a vertical axis on a table by means of a vertical handle mounted on the rim of the wheel. ‘Turn wheel in time with heard whoops’ means rotate the wheel in time with an unbroken sequence of the 1.2 s whoop sounds. ‘Sway side-to-side with imagined whoops’ means imagine whoops sounding in your head and sway in time with them.

**Fig.1.**
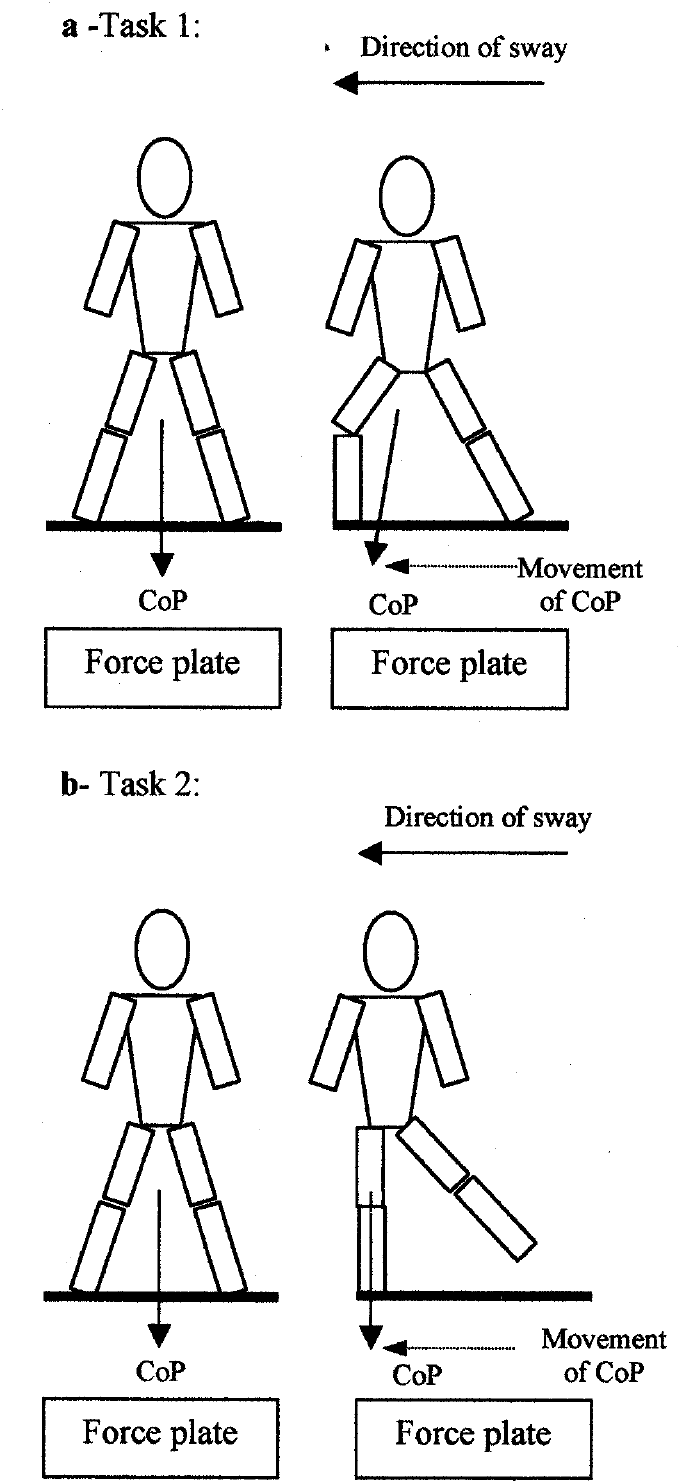
(a) Task 1: sway from side-to-side without lifting feet. (b) Task 2: sway from side-to-side lifting trailing foot.

**Table 2.** Experimental protocol.

|  |  |  |
| --- | --- | --- |
| **Session 1** |  |  |
| **C1.** Sway side to side.<br>Keep feet down. | Turn wheel. | **C2.** Sway side to side.<br>Keep feet down. |
| **C1L.** Sway side to side.<br>Lift trailing foot. | Turn wheel. | **C2L.** Sway side to side.<br>Lift trailing foot. |
| **Session 2** |  |  |
| **C3.** Sway side to side.<br>Keep feet down. | Turn wheel in time with<br>heard ‘whoop’ sounds. | **C4.** Sway side to side,<br>imagining the ‘whoop’<br>sounds. Keep feet down. |
| **C3L.** Sway side to side.<br>Lift trailing foot. | Turn wheel in time with<br>heard ‘whoop’ sounds. | **C4L.** Sway side to side<br>imagining the ‘whoop’<br>sounds. Lift trailing foot |

## Measuring *tauG*-control of body sway

The degree to which body sway was *tauG*-controlled was measured by analyzing the movement of the centre of pressure of the feet on the force plates. Since this movement was principally in the lateral *x*-direction, the analysis was based on the changing *x*-coordinate of the centre of pressure of the feet. The *x* time series, {*x*}, was analyzed using a bespoke computer program (TauServer). The program steps were as follows. (i) The 1000 Hz time series {*x*} was smoothed with a Gaussian filter, sigma 22, to reduce noise in the data. This resulted in a time series {*x_sm_*} with a cut-off frequency of 8.3 Hz. (ii) The {*x_sm_*} time series was numerically differentiated with respect to time to yield the velocity time series {*x_sm_vel*}. (iii) The peak values in the velocity time series, {*x_sm_vel*}, were found, together with the start-time, *t_start_*, and end-time, *t_end_*, of each particular sway movement. *t_start_*, and *t_end_* were defined respectively as the times before and after the peak velocity when the velocity just exceeded 10% of the peak velocity between *t_start_* and *t_end_*. This was to eliminate noisy estimates of *tau* when the velocity is low (since tau is the reciprocal of velocity). (iv) The values of *x_sm_* at times *t_start_*, and *t_end_* were recorded as *x_sm,start_* and *x_sm,end_*. (v) At each time, *t*, during each sway (for *t* running from zero to *T*, the duration of the sway), *τ_X_* was computed as (*x_sm_–x_sm,end_*)/*x_sm_vel*), and *τ_G_* was computed as 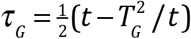, as per Eq. (1). (vi) *τ_X_* was linearly regressed recursively on *τ_G_*, to derive the following two measures of the degree to which closure of the gap *X* (= *x_sm_* – *x_sm,end_*) was *tauG*-controlled. (a) The *percentage of the gap that was tauG-guided*, namely the highest percentage of data points up to the end of the movement that fitted the *tauG*-control equation 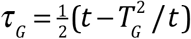, with less than 5% of the variance unaccounted for (i.e., with *r*^2^ of the linear regression greater than 0.95). (b) The *percentage of the variance explained* by the *tauG-control*, which equals 100*r*^2^. The slope of the linear regression estimates the value of the coupling constant, *k*, in the *tauG*-control equation 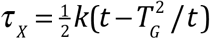.

## Results

Fig. 2 shows typical records of the displacement of the centre of pressure (CoP) when swaying side-to-side keeping feet down (Task 1), for a healthy participant (control) and an age-matched Parkinson’s patient, in baseline condition C1. Figs. 2C and 2D show the displacement and velocity of the CoP for the single sway indicated. The vertical grey lines show the start and end points for the tauG analyses of the movements of the CoP towards the left. The recursive regression analysis for the control shows strong tauG-control of body sway (percentage of movement tauG-coupled = 100%, *r*^2^ = 0.99, *k* = 0.5), whereas the tauG-control of body sway for the Parkinson’s patient is very much weaker, if it may be said to exist at all (percentage of movement tauG-coupled = 23%, *r*^2^ = 0.95, *k* = 0.8).

**Fig. 2.**
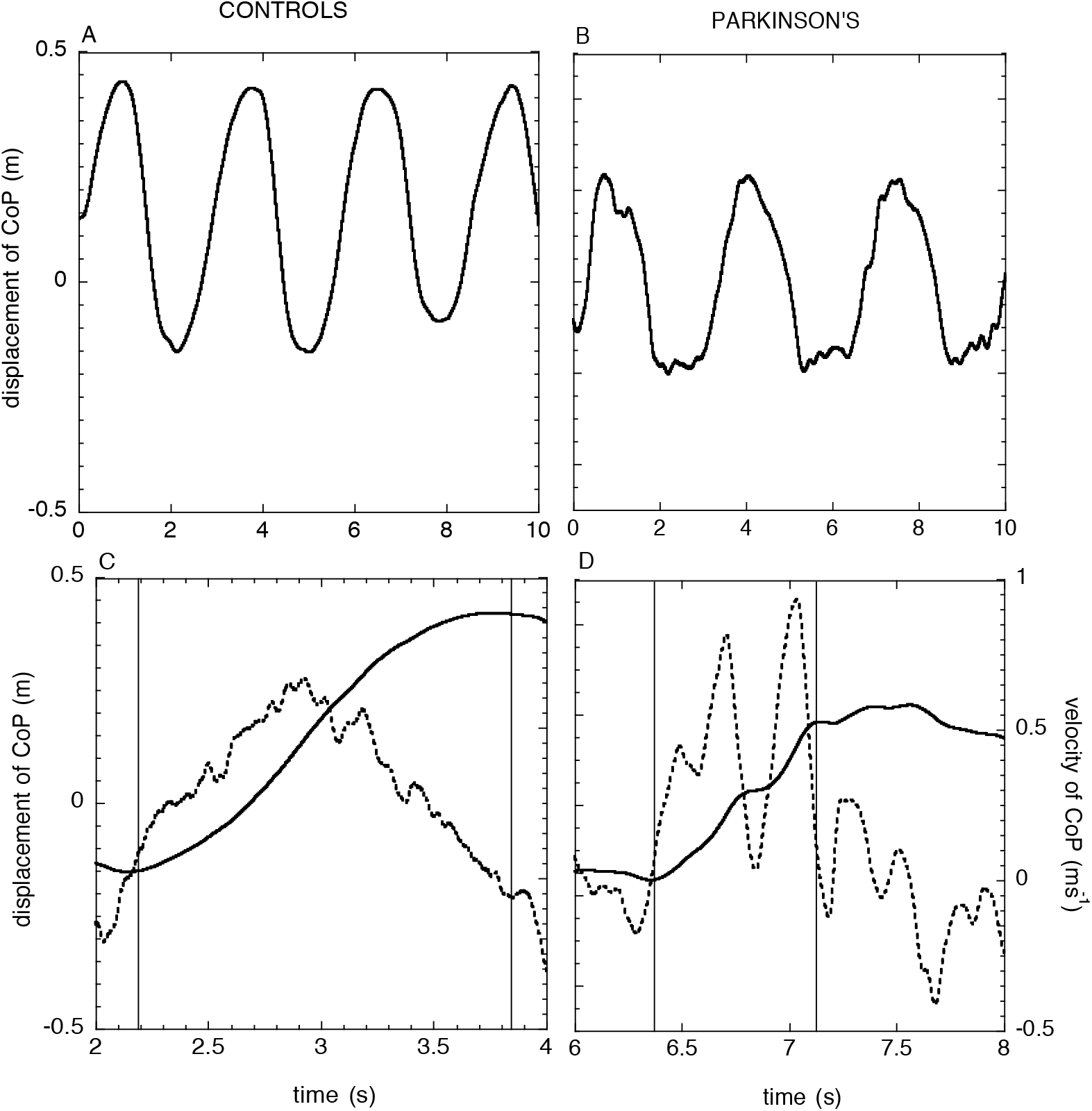
Movement of centre of pressure (CoP) when swaying side-to-side keeping feet down (Task1), in baseline condition C1. (A) Healthy participant (control), (B) age-matched Parkinson’s participant. (C) and (D) zoomed-in portions of (A) and (B), as indicated on the time axes. The broken-lines graph the velocity of the CoP. The vertical grey lines indicate the start and end points for the *tauG* analysis of the movement of the CoP to the participant’s left.

Fig. 3 shows typical records of the displacement of the centre of pressure (CoP) when swaying side-to-side lifting the trailing foot (Task 2), for a healthy participant (control) and an age-matched Parkinson’s participant, in baseline condition C1L. The leftward movement is less controlled for the Parkinson’s participant, since it has two velocity peaks.

**Fig. 3.**
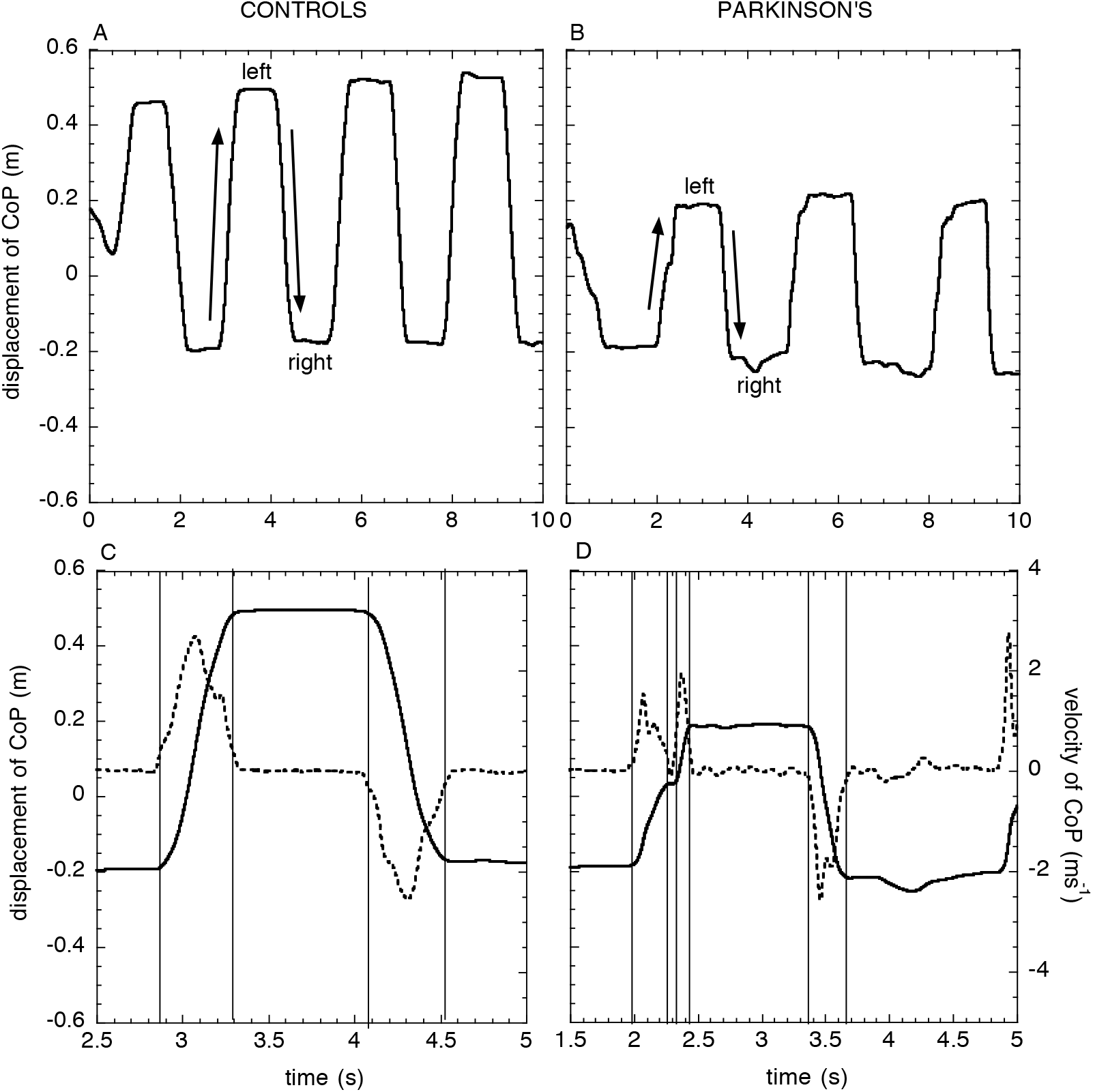
Movement of centre of pressure (CoP) when swaying side-to-side lifting trailing foot (Task 2), in baseline condition C1L. (A) Healthy participant (control), (B) age-matched Parkinson participant. (C) and (D) zoomed-in portions of (A) and (B), as indicated on the time axes. Each show a rightward sway followed by a leftward sway. The broken-lines graph the velocity of the CoP. The vertical grey lines indicate the start and end points for the *tauG* analysis of the displacement of the CoP.

To assess the effects of practice turning the wheel with and without the whoop sounds, we first computed, from the movement of the centre of pressure of the feet on the force plates (Methods), two performance measures: *% tauG-coupled*, the percentage of the sway data that fitted the *tauG* function to a criterion *r*^2^ > 0.95; and *k of tauG-coupling*, the *k* value of the tauG coupled sway movements. Performance was measured over fourteen consecutive sways (seven to the left, seven to the right) for each participant and each experimental condition. The mean % tauG-coupled and the mean k values of the tauG-couplings over each set of fourteen sway movements were then calculated.

The means are plotted in Fig. 4 for Task1 and in Fig. 5 for Task2. T1 is the baseline test; T2 is the test after turning a handle without sound; T3 is second baseline test, taken about 2 weeks after T1 and T2; T4 is the test after turning the handle accompanied with whoop sounds. Participants are colour-coded.

**Fig. 4.**
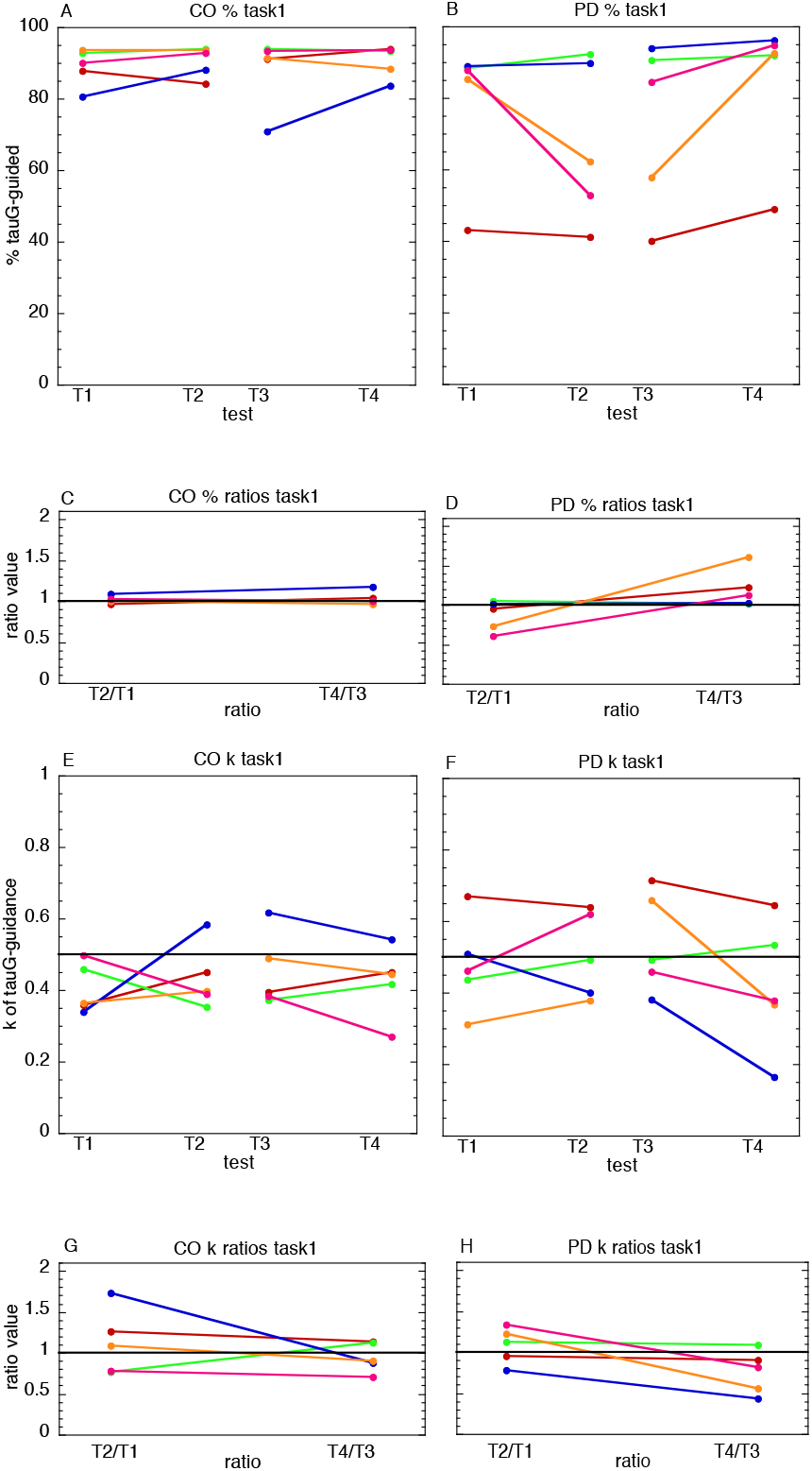

**Fig. 5.**
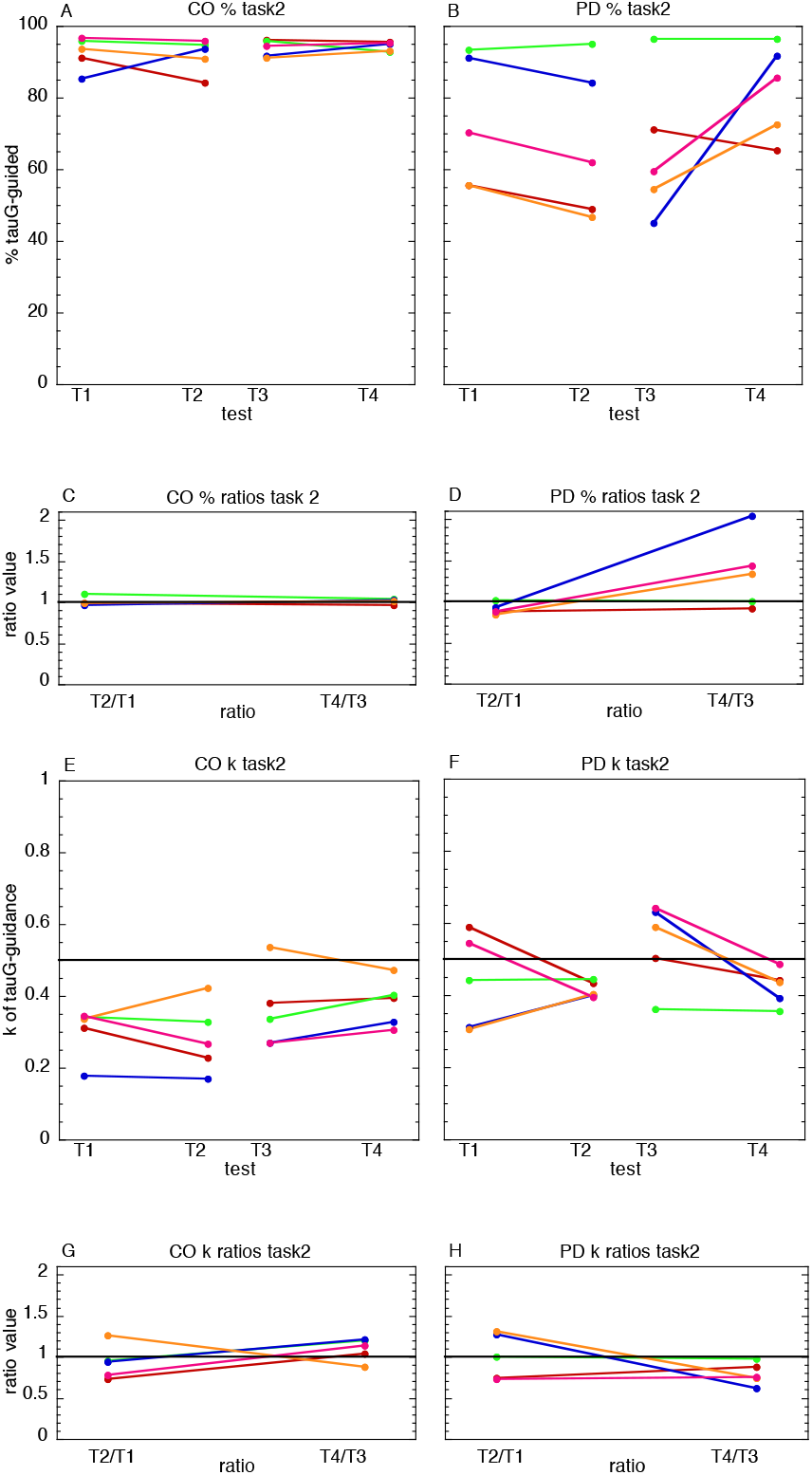

Figs. 4A and 5A show that across the 4 tests the control participants were generally quite high and consistent in their % tauG-coupling scores: average (sd) 89.55 (5.94) and 93.14 (3.41) for Task1 and Task2 respectively. In contrast, Figs. 4B and 5B show that the Parkinson’s % tauG-coupling scores were lower and less consistent: average (sd) 76.10 (0.09) and 72.01 (18.24) for Task1 and Task2.

Figs. 4E and 5E show that across the 4 tests the control participants were generally low and consistent in their k of tauG coupling scores: average (sd) 0.43 (0.09) and 0.33 ( 0.09) for Task1 and Task2 respectively. In contrast, Figs. 4F and 5F show that the Parkinson’s k of tauG-coupling scores tended to be higher and less consistent: average (sd) 0.48 (0.14) and 0.46 (0.10) for Task1 and Task2 respectively. This indicates that the Parkinson’s participants were generally slightly more abrupt in their movements than the control participants.

The smaller panels (C, D, G, H) in Figs. 4 and 5 plot the ratios of the mean scores on tests T2 and T1, and on tests T4 and T3. The ratios T2/T1 and T4/T3 equal the ratio improvements in score after turning the handle without or with whoop sounds, respectively. In the case of k of tauG-control, the ratios plotted are the reciprocals of T2/T1 and T4/T3, because a less abrupt (lower k) movement represents an improvement. Thus, in general, a graph line up to the right indicates there was a greater ratio improvement following handle-turning with whoop sounds compared to without sounds.

Table 3 shows that the number of participants (out of five) who showed greater ratio improvement following practice with whoop sounds compared to practice without sounds was generally higher for the Parkinson’s participants, for whom the number was 4 or 5.

**Table 3.** Number of participants out of five who showed greater ratio improvement following practice with whoop sounds compared to without sounds.

|  | Task1 | Task1 | Task2 | Task2 |
| --- | --- | --- | --- | --- |
|  | CO | PD | CO | PD |
| % tauG-coupling | 2 | 4 | 3 | 4 |
| k tauG-coupling | 4 | 5 | 1 | 4 |

## DISCUSSION

A theory of action control (General Tau Theory) was applied to analyzing normal and abnormal movements in PD; and to designing and testing the efficacy of a sonic aid for PD. A central aspect of the theory, which is supported by experimental evidence across a variety of actions and species, is that the trajectories of competent skilled actions follow a particular temporal pattern, which is described by the mathematical function, tauG. TauG-control of actions is deficient in PD. Therefore, we designed a device that generates whoop-like sounds, where the fundamental frequency of the sound follows the tauG pattern. Our hypothesis was that by listening to these sounds the nervous system of someone with PD might be helped subsequently to self-generate tauG patterns in their nervous system, which might facilitate movement control in different situations. Five adults with PD, and five age-matched controls took part in the study. They each listened to the sounds under two conditions: (a) experimental - turning a handle for 5 minutes while the sounds were played (b) control – turning the handle without the sounds. Before and after each condition, the tauG-control of lateral body sway while standing was measured, using force-plates, on two tasks: (i) keeping the feet down, (ii) lifting the trailing foot. The number of participants out of five, who showed greater ratio improvement following practice with whoop sounds compared to without sounds, was, on each task, higher for the PDs than the age-matched controls (4 vs 2 or 3). Thus, for the PDs, listening to the tauG whoop-like sounds while performing one action (handle turning) improved subsequent tauG-control on a different task (body-swaying).

It therefore appears that someone with PD can benefit in the tauG-control of movement by being exposed to sensory stimulation with a strong tauG component. In our experiments we used a pure tauG sound (a ‘whoop’), which appeared to stimulate the nervous system quite effectively, even with only 5 minutes exposure to the sounds. However, this might not be the most effective way of stimulating the nervous system with the tauG pattern. Another possibility is to use music, which often contains tauG-patterns (Schogler et al. 2008). Music has emotional impact too, of course. Also it can more pleasant than listening to a series of whoops! Therefore a possible way of improving on the whoops therapy would be to use music that contains a strong tauG element and which the person enjoys listening to.

